# Adaptive gene loss in the common bean pan-genome during range expansion and domestication

**DOI:** 10.1101/2023.11.23.568464

**Authors:** G Cortinovis, L. Vincenzi, R. Anderson, G. Marturano, J.I. Marsh, P.E. Bayer, L. Rocchetti, G. Frascarelli, G. Lanzavecchia, A. Pieri, A Benazzo, E. Bellucci, V. Di Vittori, L. Nanni, J.J. Ferreira Fernández, M. Rossato, O.M. Aguilar, P.L. Morrell, M. Rodriguez, T. Gioia, K. Neumann, J.C. Alvarez Diaz, A. Gratias-Weill, C. Klopp, V. Geffroy, E. Bitocchi, M. Delledonne, D. Edwards, R Papa

**Author notes:** Co-first authors (G Cortinovis, L. Vincenzi). These authors contributed equally to this work (M. Delledonne, D. Edwards, R Papa).

## Abstract

The common bean (*Phaseolus vulgaris* L.) is a crucial grain legume crop [1,2] whose life history offers an ideal evolutionary model to identify and study adaptive variants in wild and domestication populations [3]. Here we present the first common bean pan-genome based on five high-quality genomes and whole-genome reads representing 339 genotypes. We found ∼243 Mb of additional sequences containing 7,495 protein-coding genes missing from the reference, constituting 51% of the total presence/absence variations (PAVs). There were more putatively deleterious mutations in PAVs than core genes, probably reflecting the lower effective population size of PAVs as well as fitness advantages due to the purging effect of gene loss. Our results suggest strong pan-genome shrinkage occurred during wild range expansion from Mexico to South America, with more PAV loss per individual in Andean vs Mesoamerican populations. Selection signatures during wild spreading and domestication were also associated with PAV loss involved in important adaptive traits. Our findings provide evidence that partial or complete gene loss was a key adaptive trait leading to localized and genome-wide reductions. This novel result has major implications for the understanding of the process of plant adaptation and claims for a paradigm shift in evolutionary genetics. Moreover, the common bean pan-genome is a valuable resource for food legume research and breeding towards climate change mitigation, and sustainable agriculture.

## Main

Food legumes provide valuable genetic resources to address agriculture-related societal challenges, including climate change mitigation, biodiversity conservation, and the need for sustainable agriculture and healthy diets [4–7]. The common bean (*Phaseolus vulgaris* L.) is a diploid (2n=2x=22) and predominantly self-pollinating annual grain legume crop with a prominent role in agriculture and crucial societal importance [1,2]. It is also an ideal model of crop evolution [3] reflecting the parallel and independent life history of two geographically isolated and genetically differentiated gene pools (Mesoamerican and Andean) following the expansion of wild from Mexico to South America ∼150,000–200,000 years ago, an order of magnitude earlier than its dual domestication [8–11]. Previous studies using a single reference genome have provided insights into the population structure of the common bean [12] and the genetic basis of important adaptive traits [13]. However, pan-genomic diversity must be explored to gain a more comprehensive understanding [14–17]. We therefore constructed the first *P. vulgaris* pan-genome using a non-iterative approach and investigated its genetic variation in terms of PAVs within a representative panel of genetically and phenotypically well-characterized accessions. This publicly available common bean pan-genome provides a valuable starting point to identify genes and genomic mechanisms affecting adaptation and will accelerate legume improvement.

### Characterization of the common bean pan-genome

To generate the common bean pan-genome, we applied a non-iterative approach to five high-quality *de novo* genome assemblies of wild and domesticated genotypes and incorporated short-read whole genome sequencing (WGS) data from 339 representative common bean accessions, comprising 33 wild and 306 domesticated forms. This revealed ∼242 Mb of additional sequence containing 7,495 genes missing from the reference genome. The new sequences account for 22% of all discovered genes, with 9% (3,040 genes) derived from the high-quality genomes and the remaining 13% (4,455 genes) from the panel of 339 WGS genotypes. The final size of the reconstructed pan-genome was ∼780 Mb, with 34,928 predicted protein-coding genes (**Supplementary Table 1a**).

The new reference pan-genome was used for both variant calling and PAV calling (**Supplementary Table 1b**). We detected 23,343,365 variant sites, 19,002,047 of which were classified as single-nucleotide variants (SNVs) and 4,341,318 as insertions/deletions (InDels). Following PAV calling, the categorization of all 34,928 predicted genes by frequency unveiled that 58% of the pan-genome consists of core genes found across all lines (20,369 genes), with the remaining 42% comprising PAVs. These PAVs are either partially shared among accessions or exclusive to a single genotype, totalling 14,559 genes (**Supplementary Table 1c**). Notably, 51% of these PAVs originate from non-reference regions (NRRs), representing sequences absent in the reference genome. The growth curve related to the size calculation suggested a closed pan-genome. In agreement, the pan-genes reached the saturation point (99%, 34,579 genes) and remained constant without substantial increase when the number of accession genomes exceeded 120. In contrast, the size of the core genes decreased with each added genotype (**Fig. 1a**). This indicates that the final pan-genome includes almost all the gene content of *P. vulgaris*. Gene Ontology (GO) enrichment analysis showed that the core genes are significantly enriched for terms associated with homeostatic (GO:0042592) and catabolic (GO:0043632) processes (**Supplementary Fig. 1a; Supplementary Table 1d**) whereas the PAVs are significantly enriched for terms related to defence (GO:0006952), responses to external stimuli (GO:0009605), responses to light (GO:0019684), and reproduction (GO:0000003, GO:0022414) (**Supplementary Fig. 1b; Supplementary Table 1e**).

**Fig. 1:**
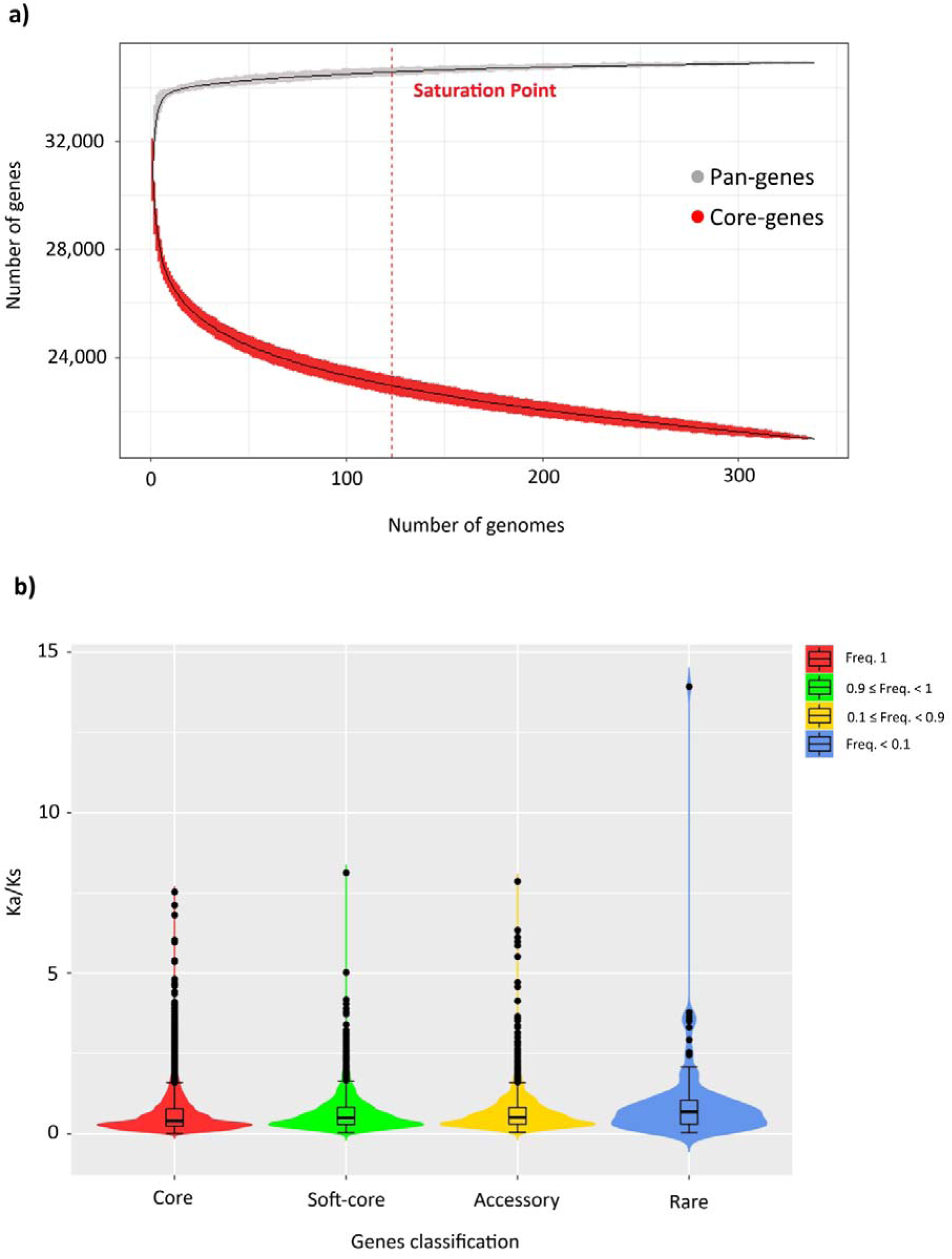
Characterization of the common bean pan-genome. **a,** Pan-gene and core gene size calculation. The growth curve of pan-genes (grey) reached saturation point (99%, 34,579 genes) when 120 individuals were included, as indicated by the dashed red line. In contrast, the growth curve of core genes (red) diminished with the addition of each genotype. **b,** Violin plots showing analysis of variance (ANOVA) related to the ratio of non-synonymous to synonymous mutations in the core genes and PAs. The PAVs are split into three subcategories based on their frequency: soft core, accessory, and rare. **Supplementary Table 1g** contains detailed statistics.

To investigate the evolution of the core genes and PAVs, we calculated the non-synonymous and synonymous ratio (Ka/Ks) for each gene in each accession (**Supplementary Table 1f**). This revealed a statistically significant difference (*p* < 2.2 × 10^−16^), with the PAVs including a greater number of harmful variants relative to benign variants when compared to the core genes (**Supplementary Fig. 1c; Supplementary Table 1g**). When we split the PAVs into three subcategories based on their frequency (*soft-core* 0.90 ≤ freq. < 1; *accessory* 0.10 ≤ freq. < 0.90; and *rare* freq. < 0.10), we observed a significant increase (*p* = 0.048) in the proportion of putative harmful variants among the rare genes compared to the soft-core genes (**Fig. 1b; Supplementary Table 1g**). These results may reflect the lower effective population size of the PAVs (reducing the efficiency of purifying selection) and/or the higher fitness gain from purging genes that have accumulated deleterious mutations (loss-of-function mutations).

### Evolutionary trajectory of the common bean

The common bean is characterized by three eco-geographic gene pools. The two major ones are the Mesoamerican (M) and Andean (A) populations, which encompass both wild and domesticated forms. The third originates from Northern Peru/Ecuador (PhI) and has a relatively narrow distribution of only wild individuals [11]. The Mesoamerican and Andean gene pools include five domesticated subgroups (M1, M2, A1, A2 and A3) corresponding to the Jalisco-Durango, Mesoamerica, Nueva Granada, Peru, and Chile races [13]. We constructed a neighbour-joining (NJ) phylogenetic tree (**Fig. 2a**) and conducted PAV-based principal component analysis (PCA) (**Fig. 2b**), both of which confirmed this well-defined population structure. Both analyses further divided the M1/Jalisco-Durango races into two clusters that we named cluster A and cluster B, respectively. The analysis of variance conducted on M1/Jalisco-Durango accessions, considering the first component for the days to flowering, revealed that cluster A is significantly later-flowering than cluster B (**Fig. 2c; Supplementary Table 2a**). The Jalisco (cluster A) and Durango (cluster B) races are therefore genetically distinguishable based on photoperiod sensitivity. This outcome also confirmed that our pan-genome enhances the characterization of genetic diversity and improves its analysis, exploitation and management. Cumulatively, the first and the second principal components of the PAV-based PCA explained 47.2% of the total variance, where PC1 mainly defined the differences between Mesoamerican and Andean gene pools while PC2 split the groups and subgroups within each gene pool (**Fig. 2b**). The NJ tree further underscored the suitability of core genes for phylogenetic reconstruction because they mitigate biases arising from the absence of genetic material among the compared accessions. In contrast to the tree based on single-nucleotide polymorphisms (SNPs) located on PAVs (**Supplementary Fig. 2a**), the NJ tree based solely on core SNPs properly grouped the wild PhI accession close to the wild Mesoamerican genotypes originating from Guatemala and Costa Rica (**Fig. 2a**), which are most closely related to the PhI gene pool [11].

**Fig. 2:**
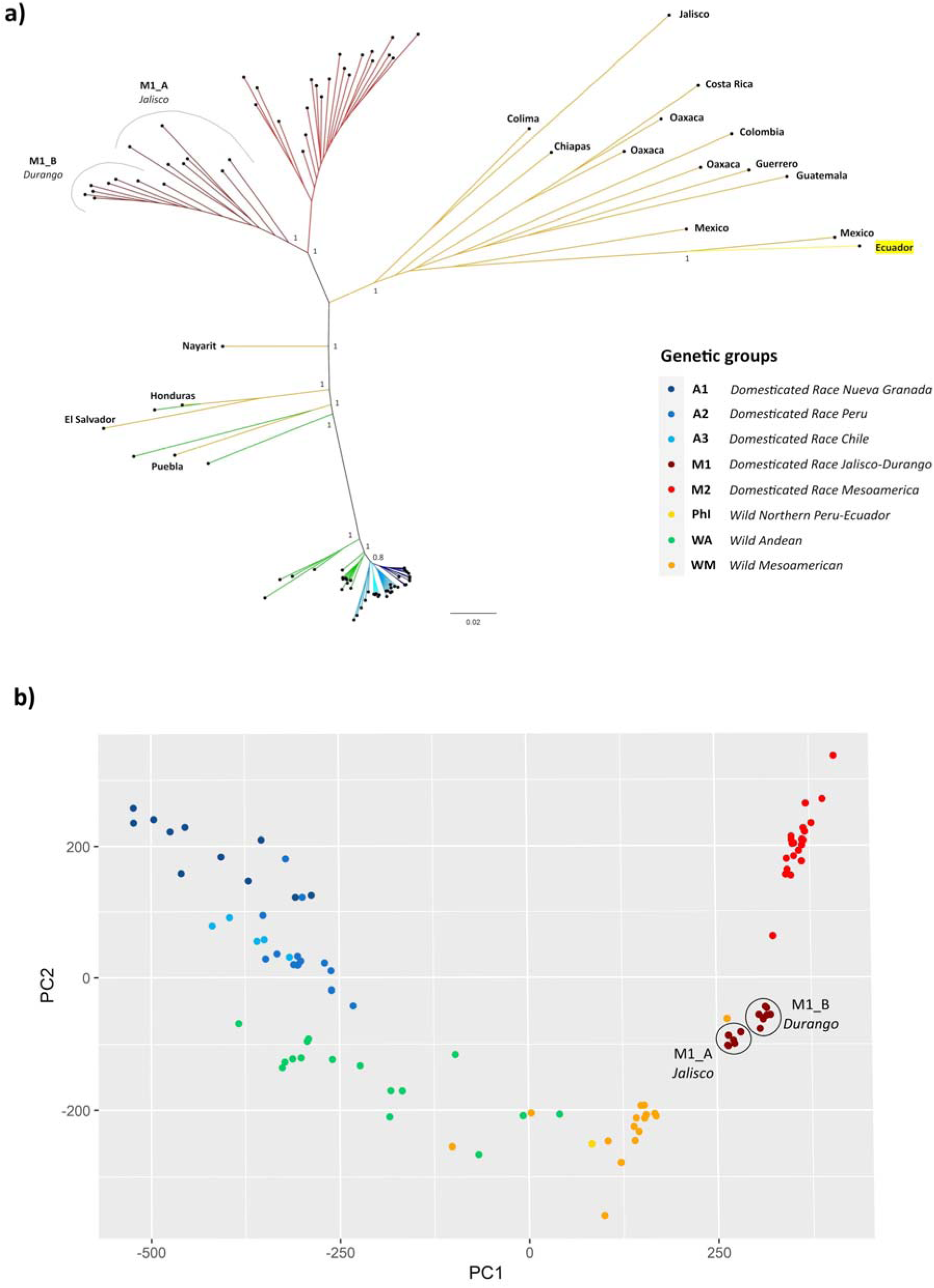
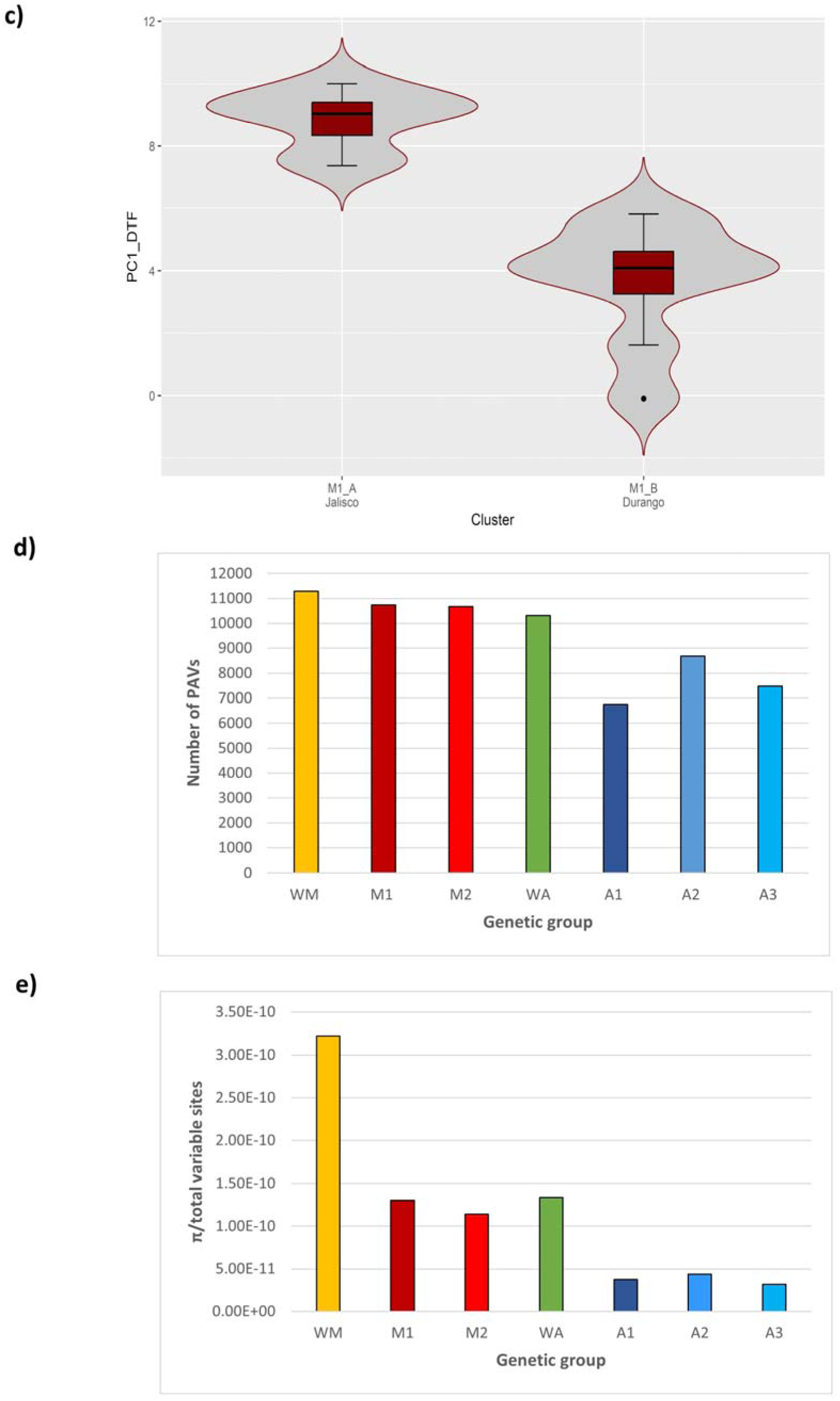
Population structure of P. vulgaris. **a,** Neighbour-joining (NJ) phylogenetic tree constructed using only SNPs located in core genes (bootstrap = 1000). **b,** PAV-based principal component analysis (PCA). **c,** Violin plots showing the analysis of variance (ANOVA) for PC1 representing days to flowering and photoperiod sensitivity in the M1/Jalisco-Durango races by splitting the accessions into two clusters based on PCA and the NJ tree. **d,** Bar chart showing the number of PAVs per genetic group. **e,** Bar chart showing nucleotide diversity calculated by estimating *π* in 250-kb windows. All procedures were applied to a representative subset of 99 genetically and phenotypically well-characterized *P. vulgaris* accessions.

When we examined the total number of PAVs per genetic group (**Supplementary Table 2b**), we found that the wild Mesoamerican and Andean populations have more PAVs compared to their domesticated counterparts (**Fig. 2d**). This supports the well-established notion that domestication is usually associated with a reduction of genetic diversity. The amplification of gene loss in domesticated common bean could reflect a classic bottleneck effect rather than natural selection [18]. We also found that the M1/Jalisco-Durango and A2/Peru races have more PAVs than the other subgroups in the same gene pool (**Fig. 2d**). This was supported by nucleotide diversity analysis applied to the 1,451,663 core SNPs (**Fig. 2e** ; **Supplementary Table 2c**) and agrees with a recent hypothesis proposing that the M1/Durango-Jalisco and A2/Peru races were the first domesticated Mesoamerican and Andean populations from which the M2, A1 and A3 races arose during a secondary domestication phase [13].

To study inter-gene-pool hybridization, the PAV matrix for American domesticated accessions was analysed by using Fisher’s exact test to compare the Mesoamerican and Andean populations. We found 5,556 PAVs (65% of the total) with a statistically significant difference in frequency (*p* < 0.05) between the two gene pools. These included 778 diagnostic PAVs, 91% (707) of which were fixed in the Mesoamerican gene pool and 9% (71) in the Andean gene pool (**Supplementary Table 2d**). GO enrichment analysis applied to the 778 diagnostic genes revealed enrichment in processes related to detoxification (GO:0098754), metabolism (GO:0008152), and responses to stimuli (GO:0050896) (**Supplementary Fig. 2b**). Interestingly, none of these PAVs were found to be diagnostic between gene pools in Europe (**Supplementary Table 2d**), and when Fisher’s exact test was applied to the subset of 114 European accessions, we did not detect any diagnostic genes between the Mesoamerican and Andean gene pools (**Supplementary Table 2e**). These outcomes clearly reflect the extensive inter-gene-pool hybridization in European germplasm and confirm its key role in the adaptation of common bean to new agricultural environments [13].

To investigate the influence of PAVs on important trait (i.e., flowering time) variations and identify candidate genes associated with them, we conducted a PAV-based genome-wide association study (GWAS) involving 218 American and European domesticated genotypes. We identified 39 significative association events correlated with day-to-flowering and photoperiod sensitivity, previously detailed in [13]. These associations were linked to 35 potential candidate PAVs, highlighting their likely involvement in regulating floral transition (**Supplementary Table 2f**). An interesting example is the GWAS peak associated with flowering time and photoperiod sensitivity located on Phvul.003G185200 (Chr03:40,838,810-40,850,729). This PAV demonstrates orthology to the *HDA5* gene in *Arabidopsis thaliana*, which displays deacetylase activity. Notably, *A. thaliana* mutants with impaired HDA5 expression patterns manifest late-flowering phenotypes due to the up-regulation of two floral repressor genes, namely FLOWERING LOCUS C (FLC) and MADS AFFECTING FLOWERING 1 (MAF1) [19]. It is noteworthy that common bean genotypes lacking the PAV Phvul.003G185200 exhibit early flowering phenotypes compared to those accessions carrying the gene (**Supplementary Fig. 2c**), implying an adaptive role correlated to the gene loss. Furthermore, we found that 9 out of the 35 candidate PAVs for GWA analysis show signature of selection in various comparisons, specifically, two PAVs between wild and domesticated Mesoamerican populations and seven PAVs between wild and domesticated Andean populations. Overall, although the majority (59%) of the candidate PAVs were located on the reference genome, a significant 41% were situated on the NRRs (**Supplementary Table 2f**), reaffirming the efficacy of the pan-genome in identifying functional variants associated with economically or evolutionarily important traits.

### Pan-genome shrinkage during wild expansion to South America

One of the most striking outcomes we observed was the difference in pan-genome size between the Mesoamerican and Andean gene pools (**Fig. 3a**). We calculated the total number of PAVs per individual and found that accessions from the same gene pool clustered together in separate groups, with Mesoamerican accessions featuring more PAVs per accession than Andean ones (**Fig. 3b, c; Supplementary Table 3a**). One possible explanation is that this reduction in pan-genome size may simply reflect genetic drift and the two sequential bottlenecks that occurred solely in the Andean population [12]. To better understand the roles of different evolutionary forces in shaping the PAV content of the Mesoamerican and Andean gene pools, and to distinguish between the effects of adaptation, population demography and history, we first focused on the analysis of wild accessions. Considering a panel of wild genotypes representing the entire geographical distribution in Latin America, we applied bivariate fit analysis and found a significant correlation (*p* < 0.0001) between the number of PAVs per individual and the latitude. Analysis of variance in which wild individuals were grouped by latitude followed by spatial interpolation revealed a significant and progressive loss of genes ranging from the accessions of Northern Mexico to those of Northwestern Argentina (**Fig. 4a, b; Supplementary Table 3b**). Furthermore, F_ST_ analysis on PAVs comparing Mesoamerican and Andean wild populations may suggests the occurrence of selection for gene loss during wild range expansion (**Fig. 4c**). We found that 64% of all candidate PAVs in the top 5% of the F_ST_ distribution (Fst≥0.85) are absent in the wild Andean gene pool. This high rate of gene loss in the Andean population significantly exceeds the gene loss rate observed in the entire variable genome (26%), demonstrating a more than twofold increase. This difference in gene loss rates was statistically validated using bootstrap resampling, strongly suggesting that gene loss during the process of wild differentiation was not a random occurrence but the evident outcome of selective forces (**Supplementary Fig. 3a, b**). Moreover, functional annotation of the candidate PAVs revealed the enrichment of genes involved in pollen germination, innate immunity, abiotic stress tolerance, and root hair growth, indicating a potential adaptive role during wild range expansion (**Supplementary Table 3c**). Overall, our findings suggest that selective pressure favouring the loss of genes involved in adaptive mechanisms, coupled with the influence of genetic drift resulting from the founder effect, may have contributed to the shrinking of the Andean pan-genome during wild differentiation.

**Fig. 3:**
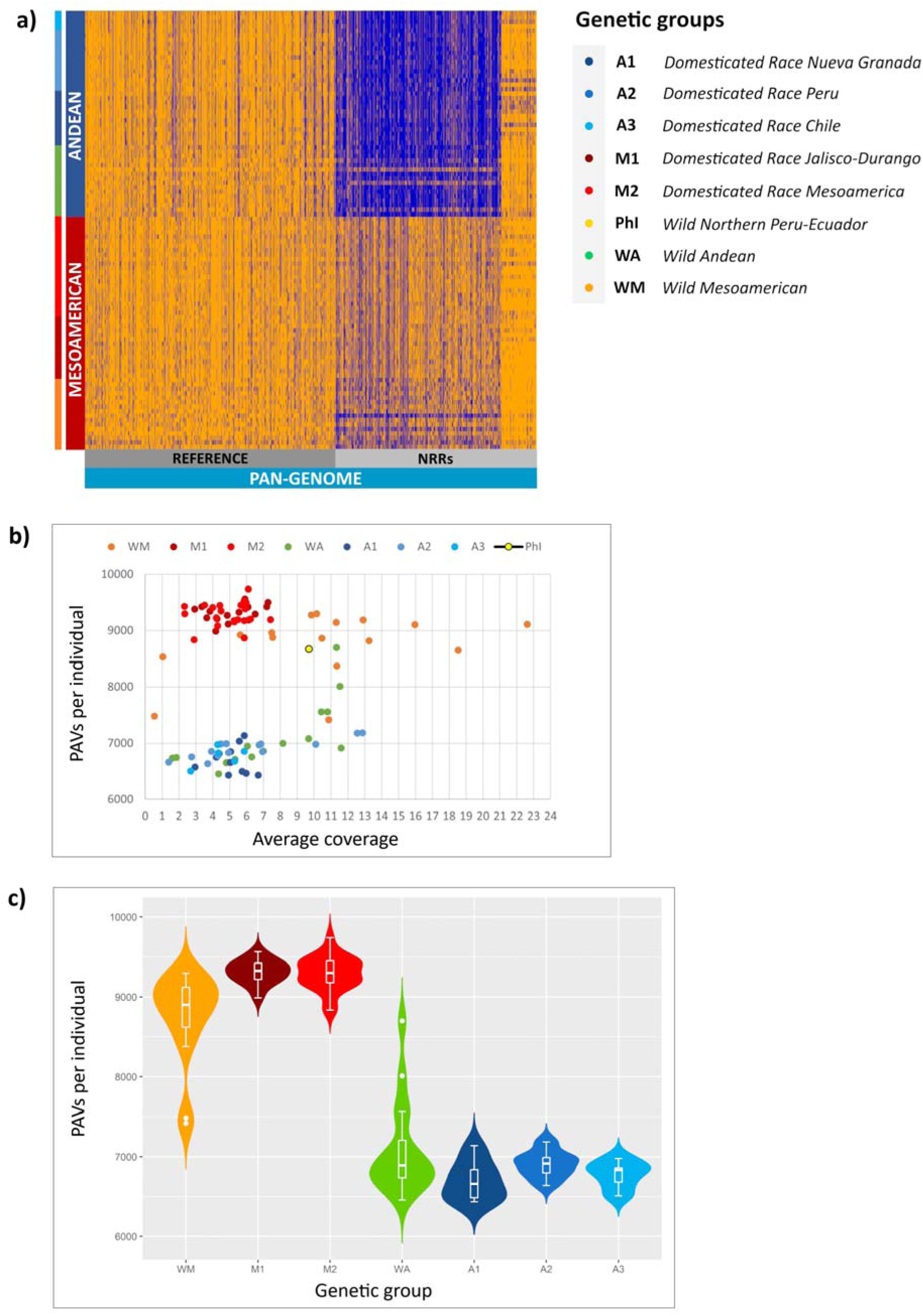
Evolution of the common bean pan-genome. **a,** Heat map illustrating the number of PAVs per individual in the final pan-genome. Orange indicates presence while blue indicates absence. **b,** Scatterplot showing the number of PAVs per individual (y-axis) in relation to the coverage (x-axis) of each genotype. **c,** Violin plots representing the analysis of variance (ANOVA) for the number of PAVs per individual by genetic group. All procedures were applied to a representative subset of 99 genetically and phenotypically well-characterized *P. vulgaris* accessions. **Supplementary Table 3a** contains detailed statistics.

**Fig. 4:**
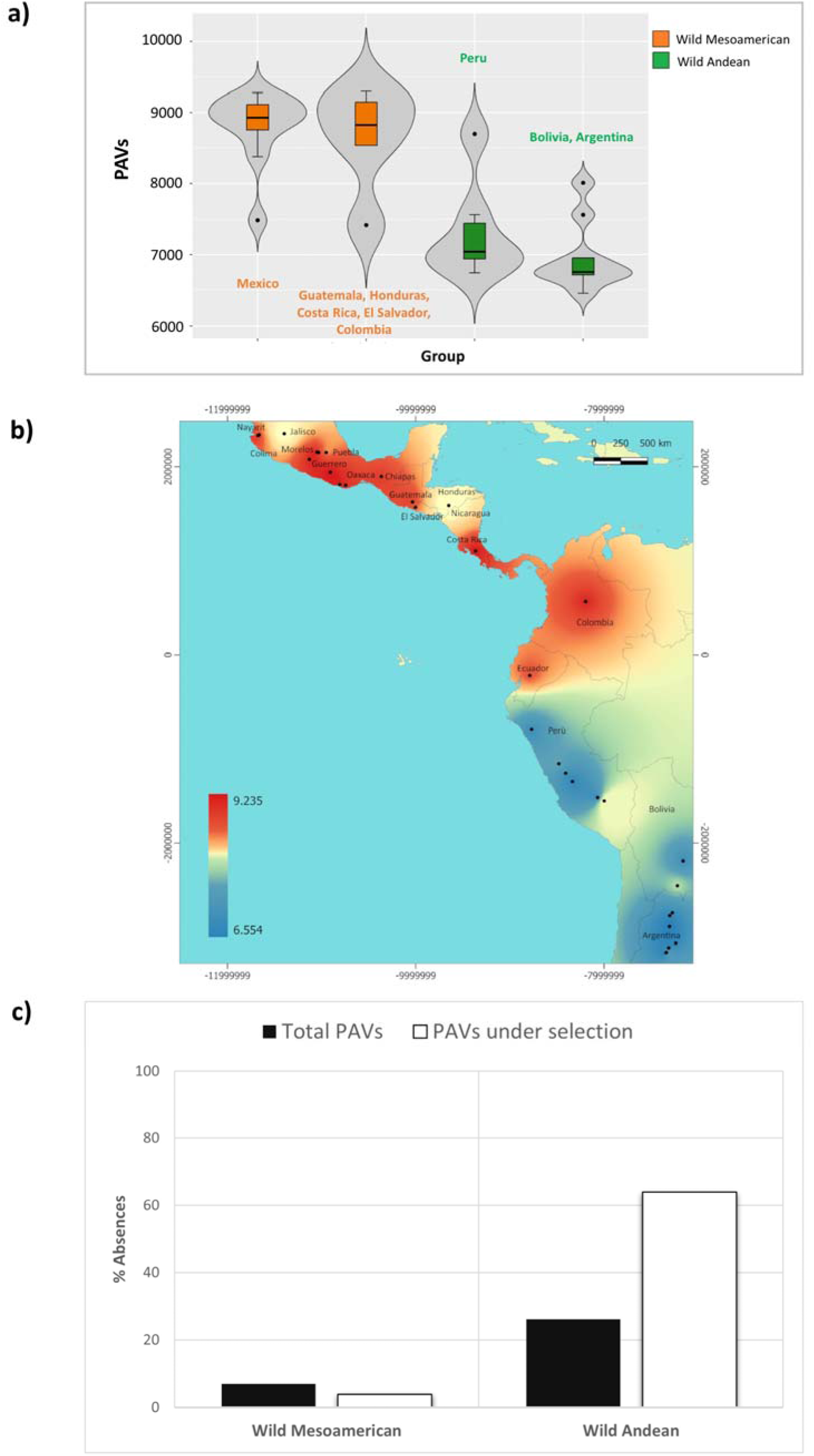
Selection for adaptive gene loss during the expansion of wild common bean. **a,** Violin plots showing the analysis of variance (ANOVA) for the number of PAVs per individual based on grouping the wild Mesoamerican and Andean accessions according to latitude coordinates. Supplementary Table 3b contains detailed statistics. b, Interpolation of the geographic distributions of the wild accessions based on the number of PAVs per individual. Dark red regions indicate a higher number of PAVs and blue regions a lower number of PAVs. c, Bar charts showing the proportions of absences found for the subset of PAVs putatively under selection during the wild expansion (white) and for the entire variable genome (black).

### Footprints of selection for gene loss during domestication

The PAVs putatively shaped by selection during domestication in Mesoamerica and the Andes revealed further evidence that gene loss underpinned the successful adaptation of the American common bean. F_ST_ analysis was applied to PAVs in wild and domesticated forms (separately for each gene pool) with only PAVs in the top 5% of the F_ST_ distribution considered as candidates. We found 460 PAVs potentially under selection in the Mesoamerican population (F_ST_ ≥ 0.31) and 514 in the Andean population (F_ST_ ≥ 0.27) (**Supplementary Table 4a, b**). Functional annotation of the candidate PAVs revealed the enrichment of genes associated with domestication syndrome and adaptive traits such as dormancy, floral transition, light acclimation, defence, and symbiotic interactions (**Supplementary Table 4c, d**). Importantly, the candidate Phvul.003G265200 (Chr03: 50,365,995-50,368,501) is orthologous to 11 members of the plant Rho GTPase subfamily (ROP), including *ROP6* encoding a small Rho-like GTP binding protein. This GTPase subfamily is required for symbiotic interactions [20–22], and in the plasma membrane of *Lotus japonicus* cells it interacts directly with NOD FACTOR RECEPTOR 5, one of two nodulation factor receptors essential for nodule formation during symbiosis [23]. From our analysis, Phvul.003G265200 is a putative selected PAV (F_ST_ = 0.50) for the Mesoamerican gene pool whose frequency declined by more than 60% during progression from the wild (0.94) to the domesticated (0.25) population (**Supplementary Table 4a**). Overall, no significant differences were observed in terms of absences between the wild and domesticated populations of both gene pools. However, a significant proportion of PAVs putatively under selection, specifically 63% (289 genes) in the Mesoamerican population (**Fig. 5a**) and 80% (411 genes) in the Andean one (**Fig. 5b**), were less frequent in domesticated than wild populations. When considering all PAVs, the percentage of PAVs with lower frequencies in domesticated populations fell significantly to 22% (*p* < 2.2 × 10^−16^) for the Mesoamerican gene pool and 29% (*p* < 2.2 × 10^−16^) for the Andean one (**Fig. 5a, b**). These findings suggest that selection during domestication resulted in gene loss, but unlike the range expansion of wild populations, it did not completely abolish the selected genes. This may reflect the different evolutionary timescales involved: wild differentiation occurred ∼150,000 years ago whereas domestication was much more recent at ∼8,000 years ago. These findings are consistent with previous observations that selection during the domestication of common bean in Mesoamerica has directly affected the transcriptome, leading to a ∼20% decrease in gene expression levels attributed to loss-of-function mutations [18]. We also detected 29 PAVs with high F_ST_ values in common between the Mesoamerican and the Andean gene pools, and these are mainly associated with the tryptophan metabolic pathway. Tryptophan holds significance as a precursor in secondary metabolism, contributing to the synthesis of essential molecules like auxin, serotonin, and melatonin. These compounds play diverse roles in plant physiology, influencing processes such as seed germination, root development, senescence, and flowering. Additionally, they contribute to the plant’s response mechanisms against both biotic and abiotic stresses [24]. We analysed their frequencies and found that ∼86% in both gene pools decreased in frequency during the progression from wild to domesticated accessions (**Supplementary Table 4e**). This may indicate a pattern of genomic convergence for some key adaptive genes between the Mesoamerican and Andean populations during their parallel domestication events.

**Fig. 5:**
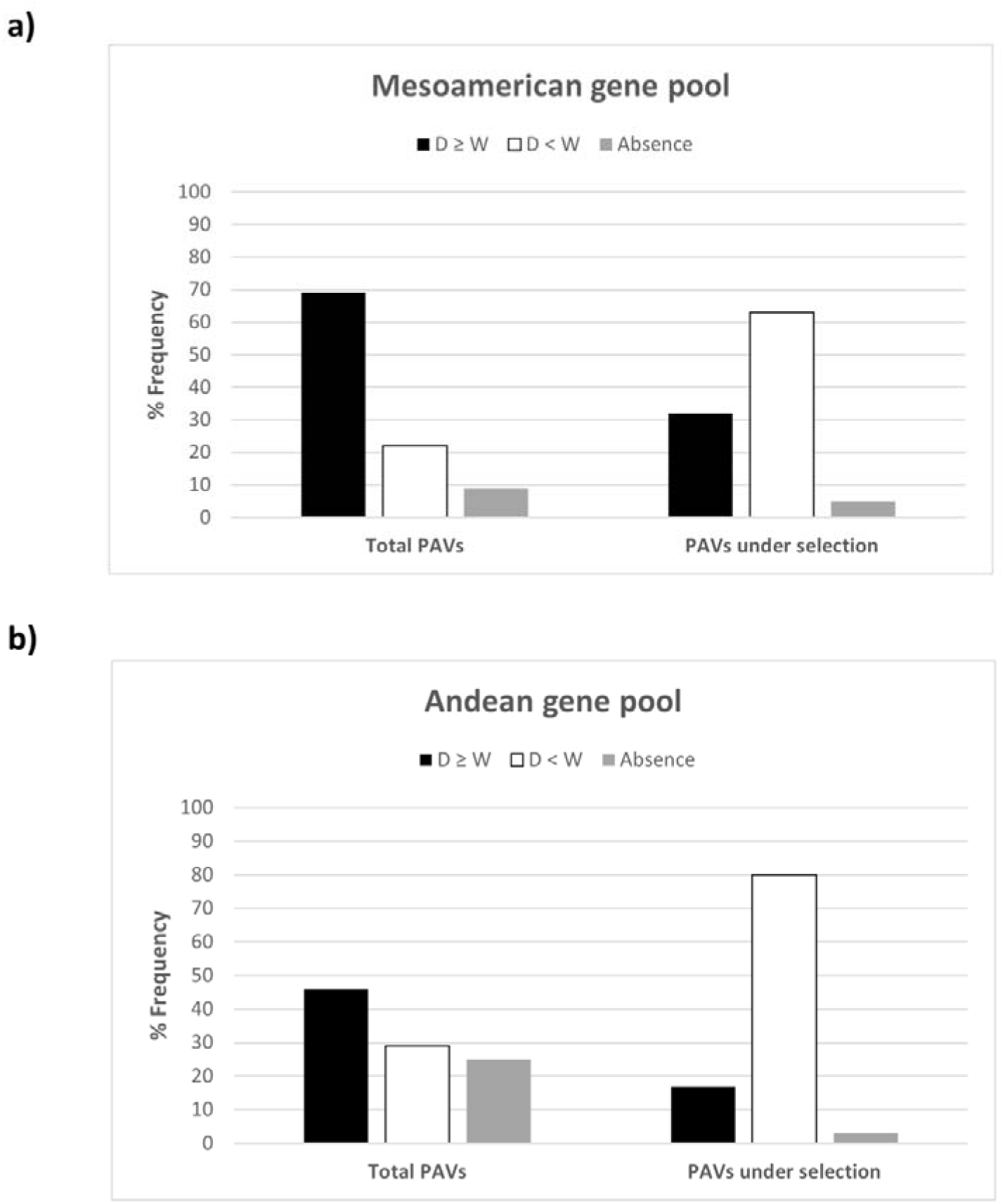
Localized adaptive reduction effects during the domestication of the common bean. **a,** Bar chart showing the proportions of presence/absence in the Mesoamerican gene pool for the entire variable genome (left) and for the subset of PAVs putatively under selection between wild and domesticated populations (right). **b,** Bar chart showing the proportions of presence/absence in the Andean gene pool for the entire variable genome (left) and for the subset of PAVs putatively under selection between wild and domesticated populations (right). In both charts, the presence values are divided based on frequency (≥/<) in the comparison between wild and domesticated forms.

## Conclusions

The global economic and social importance of the common bean means that pan-genomic analysis could boost the conservation and exploitation of its genetic resources to address key challenges in sustainable agriculture and wider society [1–6]. The genotypes selected for this study encompass both wild and domesticated forms, ensuring that the pan-genome comprehensively captures the extensive genetic variation within this species. This publicly accessible tool serves as a valuable resource for studies in population genetics, functional genomics, and plant breeding. PAV analysis provided insight into the evolutionary dynamics of pan-genome adaptation, including signals of selection for complete gene loss during wild differentiation between the Mesoamerican and Andean gene pools, contributing to the smaller pan-genome of the Andean population. We also identified selection footprints for gene loss during both domestication processes in Mesoamerica and the Andes, causing localized reductions in gene frequencies in domesticated populations compared to their wild counterparts. Interestingly, candidate genes that have been completely or partially lost appear to be involved in important adaptive mechanisms, such as flowering time, symbiosis, biotic and abiotic stress tolerance, and root hair growth.

Gene loss is considered functionally equivalent to other loss-of-function mutations, such as premature stop codons, providing an important source of adaptive phenotypic diversity [25–30]. Moreover, the variation in genome size between different populations of microorganisms and plants has been already described [31–33]. For instance, in contrast to their European native counterparts, invasive genotypes exhibited a reduced genome size resulting in phenotypic effects that could enhance the species’ invasive potential [34]. Similarly, Díez et al. [35] documented the genome size variations within the *Zea mays* species during the post-domestication process, revealing that maize landraces have significantly smaller genome sizes compared to their closest wild relatives, the teosintes. However, it is still an open question as to whether genome size variation is shaped by natural selection. Here, our results claim that under the influence of specific and diverse agro-ecological pressures, the relinquishment of particular genes can confer a selective advantage on a given population. This may be relevant when populations face selective pressure resulting from radical environmental changes, such as the expansion of wild common bean from the warm and humid climate of Central Mexico to higher and cooler altitudes in the Andes. Our research presents pioneering evidence, establishing a new paradigm in which natural selection drives gene loss, favouring adaptation over mere stochastic responses. Mutations are more likely to cause a loss rather than a gain of function, so adaptive gene loss provides a rapid evolutionary response to environmental changes. This could have profound implications for our understanding of crop adaptation under climate change. The common bean pan-genome is a valuable starting point that will lead to a deeper understanding of the genetic variations and genome dynamics responsible for key adaptive traits in food legumes.

## Methods

### Sources of genetic diversity

The pan-genome was constructed from five high-quality genomes representing the Mesoamerican and Andean gene pools. The *P. vulgaris* reference genome Pvulgaris_442_v2.0 (PV442) was downloaded from Phytozome (https://phytozome-next.jgi.doe.gov/), the genomes of BAT93 and JaloEPP558 were provided by the INRAE Institute, and the genomes of MIDAS and G12873 were sequenced and assembled *de novo* specifically for this study (**Supplementary Table 5a**). We also integrated 339 representative low-coverage WGS common bean genotypes, including 220 domesticated and 10 wild accessions from previous studies [11, 13]. The remaining 109 accessions were multiplied in the greenhouse, and DNA extracted from young leaves was used for sequencing (**Supplementary Table 5b**).

### Plant growth and DNA extraction

MIDAS and G12873 single seed descent (SSD) genotypes were multiplied in the greenhouse. For both samples, 2 g of young leaf material was collected after 48 h of dark treatment and high-molecular-weight (HMW) DNA was extracted as previously described [36]. DNA quality was evaluated according to Oxford Nanopore Technologies (ONT) requirements. Specifically, purity was assessed using a NanoDrop 1000 spectrophotometer (Thermo Fisher Scientific), the concentration was determined using a dsDNA Broad Range Assay Kit with Qubit 4.0 (Thermo Fisher Scientific), and the fragment size (≤ 400 kb) was determined using the CHEF Mapper electrophoresis system (Bio-Rad Laboratories). Fragments < 25 kb were removed using the Short Reads Eliminator kit (Circulomics) leaving 75% of the DNA from the MIDAS samples and 95% from the G12873 samples.

*P. vulgaris* genotypes of BAT93 and JaloEEP558 were sowed in soil and grown in a growth chamber at 23°C, 75% humidity, and 8 h dark and 16 h light photoperiods under fluorescent tubes (166lE). Young trifoliate leaves of BAT93 and JaloEEP558 genotypes were collected and flash-frozen with liquid nitrogen. Three days before sampling, plants were dark treated to optimize the high molecular weight DNA extraction. In addition, 109 SSD accessions were multiplied in the greenhouse and young leaves were collected in silica gel for drying and subsequent DNA extraction using the DNeasy 96 Plant kit (Qiagen) according to the manufacturer’s instructions. For each sample, 50–70 mg of dried leaf material was pulverized with a Tissue-Lyser II (Qiagen) at 30 Hz for 6 min. The DNA quality and quantity were evaluated using a NanoPhotometer NP80 (Implen), and the concentration was determined using a Qubit BR dsDNA assay kit (Thermo Fisher Scientific).

### Sequencing low-coverage WGS accessions

DNA libraries for all samples were prepared using a KAPA Hyper Prep kit and PCR-free protocol (Roche). For each genotype, 200 ng of DNA was fragmented by sonication using a Covaris S220 device (Covaris) and WGS DNA libraries were generated using a 0.7–0.8× ratio of AMPureXP beads for final size selection. Libraries were quantified using the Qubit BR dsDNA assay kit and equimolar pools were quantified by real-time PCR against a standard curve using the KAPA Library Quantification Kit (Kapa Biosystems). Libraries were sequenced on the NovaSeq 6000 Illumina platform in 150PE mode, producing 15–30 million fragments per sample.

### Sequencing and assembly of MIDAS and G12873 genomes

Following quality control and priming according to ONT specifications, libraries were sequenced on a MinION device with a SpotON flow cell (FLO-MIN106 R9.4.1-Rev D). Two libraries were prepared for each genotype according to the SQK-LSK109 ligation sequencing protocol (ONT) with minor adjustments. Each library was loaded twice, and the flow cell was washed using the Flow Cell Wash Kit (ONT). Illumina PCR-free libraries were prepared starting with 1 ug of fragmented gDNA using the KAPA Hyper prep protocol. This process involved extending the adapter ligation time up to 30 minutes and conducting post-clean-up size selection using 0.7X AMPure XP beads. The library’s concentration and size distribution were assessed using a Bioanalyzer 2100 in combination with high-sensitivity DNA reagents and chips. Sequencing was performed on a NovaSeq6000 instrument to generate 150-bp paired-end reads. MIDAS and G12873 whole-genome assemblies were generated using the nanopore-based approach based on 26 Gb (50-fold coverage) and 36 Gb (69-fold coverage), respectively. Raw nanopore reads were corrected using Canu v2.1 [37] and the resulting corrected reads were assembled *de novo* using wtdbg2 v2.5 [38]. Draft assemblies were refined by iterative polishing using long reads (Racon v1.4.3 and Medaka v1.0.3) [39] and short reads (three rounds of Pilon v1.23) [40]. Completeness was assessed using BUSCO v4.1.2 [41] and the Fabales_odb10 dataset (**Supplementary Table 5c**).

### Sequencing and assembly of BAT93 and JaloEEP558 genomes

High molecular weight DNA of BAT93 and JaloEEP558 genotypes was sequenced with a PacBio Sequel II system by GENTYANE platform (INRAE Clermont-Ferrand, France). A total of 21.09 and 29.35 Gb of PacBio HiFi reads were generated from BAT93 and JaloEEPP558, respectively. PacBio HiFi reads were *de novo* assembled into contigs using HiFiasm (v 0.9.0) with default parameters [42].

### Orthologous/paralogous analysis and clustering threshold settings

To incorporate two distinct populations, namely the Andean and the Mesoamerican gene pools, into the pangenome, precise differentiation between orthologous and paralogous genes is imperative. Consequently, a meticulous strategy was devised to ensure the preservation of solitary orthologous gene copies along with all paralogous counterparts. The relationship between orthologous genes was calculated with minimap2 v2.17 [43] to align the MIDAS and G12873 genome assemblies using the open reading frames (ORFs) of 2,330 complete single-copy BUSCO (Benchmarking Universal Single-Copy Orthologs) genes selected from the *P. vulgaris* reference genome PV442 (**Supplementary Table 5d**). The percentage identity was calculated for each ORF based on the number of matches in the alignments as a proportion of ORF length. The relationship between paralogous genes was calculated using the three most abundant gene families (OG0000273, OG0000328, and OG0000085) in the *P. vulgaris* PV442 reference genome, composed of 26, 37, and 42 genes, respectively. An all-versus-all comparison between the members of the same family was implemented using minimap2. The percentage of 1identity was calculated for each gene family by dividing the number of matches in the alignments by the reference gene ORF length and then averaging the identity percentages for each family. Finally, the results of both tests were used to establish a clustering threshold of 90% to retain only one orthologous and all paralogous genes in the pan-genome (**Supplementary Table 5e**).

### Pan-genome construction

We used a paired genome alignment strategy for pan-genome construction [44]. The PV442 reference genome was independently mapped onto MIDAS, G12873, BAT93 and JaloEPP558 with minimap2 v2.17 using the alignment pre-set *-x asm5*, which considers regions with an average divergence < 5%. The bam files of the four alignments were converted to delta files and structural variants were called using Assemblytics v 1.2.1 [45]. Only deletions were selected as NRRs [44]. Uncovered contigs private to the four analysed genomes were identified by applying samtools depth v1.1 [46] to the bam files and were extracted as NRRs. Deletions and uncovered contigs were independently filtered for a minimum length of 1 kb and clustered using CD-HIT-EST v4.8.1 [47] with a sequence identity of 90%, as described above for the orthologous and paralogous genes. To ensure that the NRRs identified through this method didn’t encompass orthologous genes already existing in the PV442 reference genome, we specifically employed highly conserved BUSCO genes. We conducted a comparative analysis between the full complement of 4,947 MIDAS and 4,812 G12873 BUSCO genes present in PV442 and the NRRs derived from MIDAS and G12873 using BLASTp. Illumina data representing the 339 low-coverage WGS common bean accessions were trimmed with fastp v0.21.0 [48] and aligned to the preliminary pan-genome using bowtie2 v2.3.5.1 [49] with default parameters. The unmapped reads from these alignments were extracted using samtools v1.11, assembled using MaSuRCA v3.4.2 [50] with default parameters, and clustered using CD-HIT-EST v4.8.1 with a sequence identity of 90%. Finally, the NRRs derived from the panel of 339 common bean accessions were added to the preliminary pan-genome to generate the final pan-genome. To exclude putative contaminants and/or organelle sequences, NRRs were compared to the NCBI non-redundant nucleotide database using BLASTn, considering a minimum 80% identity and 25% coverage, leading to the removal of 1,194 sequences. Overall, we identified 64,174 added sequences, 86% of which reflected the mapping of the 339 low-coverage WGS accessions. The remaining 14% was identified by comparing the reference genome independently with the other four high-quality genomes (**Supplementary Table 1a**).

### Pan-genome annotation

Repetitive sequences were identified and soft-masked using RepeatModeler v2.0.2 [51] and RepeatMasker v4.1.2-p1 [52]. Protein-coding genes were identified using a hybrid-approach prediction with Augustus v3.3.3 [53]. Proteins from *P. vulgaris* and correlated species (*Medicago truncatula* and *Glycine soja*) plus RNA-Seq data (unpublished data from [18]) were aligned to the genome and used as extrinsic evidence. Protein sequences were aligned with Hisat2 v2.2.1 [54] and RNA-Seq data were aligned using Genome Threader v1.7.1 [55]. BUSCO genes in the Fabales_odb10 database were used to train the model for the Augustus predictor. Predicted genes were scanned with InterProScan v5.46-81.0 [56] for the presence of protein domains. Using a custom script, genes with transposon-related domains were filtered out and retained in the annotation if they contained at least one known protein domain. The filtered proteins were compared to the pan-genome with BLASTp v2.12.0 [57] and filtered by the best hits. The predicted genes were clustered with the proteins of all species considered in the annotation using OrthoFinder v2.5.4 [58]. Finally, functional annotation was achieved by integrating information about orthologous genes and by identifying functional domains using a custom script.

### PAV calling

Illumina data representing the 339 low-coverage WGS accessions were aligned to the pan-genome using bowtie2 v2.3.5.1 and the coverage of each predicted gene was calculated for each accession using samtools v1.11 (**Supplementary Table 5f**). PAV calling thresholds were defined for each accession according to the minimum coverage value of 1000 randomly selected BUSCO genes (ORFs). The BUSCO genes are orthologous genes that should be present in all considered accessions, but a few underrepresented genes in a given accession could constitute a bias. To avoid this, values below 1% (the 10 least covered genes) were discarded. The identified genes were classified based on their frequency as core genes if present in all the accessions or PAVs if partially shared or private to a single genotype (**Supplementary Table 1b, c**).

### Pan-genes and core genes size calculation

The curves describing the pan-genome and core genome sizes were evaluated by considering 1,000 random orders of the 339 genotypes. The orders were chosen randomly among all possible permutations (n! where n=[1,339]). For each ordering, the gene sets of the accessions were progressively added to the total genome size without considering the genes already present in the total set. The same procedure was applied for the core genome size, but the gene sets were intersected when each genome was added, thus keeping only the genes in common for each iteration (**Supplementary Table 5g, h**).

### Variant calling

SNVs and InDels were called with bcftools v1.10.2 [59] based on the alignment of 339 accessions with the pan-genome using bowtie2 v2.3.5.1. We used bcftools mpileup v1.10.2 to generate a genotype likelihood table, and variants were identified using bcftools call v1.10.2 and the pileup table, producing the final vcf file.

### Non-synonymous and synonymous mutations

The Ka/Ks ratio was computed for each gene in each accession using KaKs calculator v2.0 [60]. For each gene, the consensus sequence of each accession was extracted using bcftools consensus v1.10.2. The calculator compares the pan-genome gene sequence with the gene sequence of each accession to identify non-synonymous and synonymous variants and then computes the ratio. The calculator reported *NA* when there were no variants in a specific accession or when the denominator of the Ka/Ks ratio was zero. It was possible to compute the analysis for 30,850 of 34,928 genes. Sometimes the length of one of the two compared sequences was not divisible by three so the sequence could not be read in triplets (**Supplementary Table 1f**). The average Ka/Ks value per gene was used to assess the significance of the sample median (**Supplementary Table 1g**).

## Data analysis

Pan-genome analysis focused on a representative subset of 99 well-characterized accessions among the original 339, including wild and American domesticated forms. In some cases, we also analysed the subset of 114 European domesticated accessions (**Supplementary Table 5b**).

For GO enrichment, the annotated core genes and PAVs in the pan-genome were analysed using the *buildGOmap* R function to infer indirect annotations and generate data suitable for *clusterProfiler* [61, 62]. Diagnostic genes were analysed using Metascape [63]. *A. thaliana* orthologs were identified using OrthoFinder [58] and by comparing all protein sequences from *P. vulgaris* (v2.1) and *A. thaliana* (TAIR10). For PCA, the PAV matrix (1/0) was analysed using the *logisticPCA* package in R [64].

ANOVA within subgroup M1 was carried out using the first principal component related to days-to-flowering and photoperiod sensitivity [13] as a representative phenotypic trait.

F_ST_ analysis involved the separate testing of PAVs in the Mesoamerican and Andean gene pools by comparing the frequency of each PAV between wild and domesticated forms. Each PAV was considered as a single locus (0/1) and F_ST_ was calculated by applying the formula F_ST_ = (H total – H within) / H total, where H is the heterozygosity [65]. The same procedure was applied to wild accessions when comparing the Mesoamerican and Andean gene pools. Only PAVs in the top 5% of the F_ST_ distribution were considered as candidates.

The functions of interesting PAVs and those associated with *A. thaliana* orthologs detected by OrthoFinder [58] were investigated manually in the NCBI database (https://www.ncbi.nlm.nih.gov/).

Phylogenetic analysis involved the extraction and filtering of SNVs located in core genes and PAVs using bcftools [59], resulting in two final datasets: 1,451,663 SNPs for the core genes and 338,475 SNPs for the PAVs. The datasets were used to calculate the genetic distance between individuals and compute maximum composite likelihood values with 1000 bootstraps for the NJ tree in MEGA11 [66]. The final trees were visualized in FigTree (http://tree.bio.ed.ac.uk/software/figtree/).

The filtered dataset of SNPs in core genes was also used to quantify the genetic diversity within groups of accessions by estimating *π*. The *--window-pi* vcftools flag was used to obtain measures of nucleotide diversity in 250-kb windows. The windowed-*pi* estimates were then divided by the total number of SNPs to calculate a global estimate for each genetic group.

Fisher’s exact test with the false discovery rate corrected for multiple comparisons was applied in R to identify PAVs that differed significantly in frequency between the Mesoamerican and Andean gene pools for the American and European accessions.

The phenotypic data used for PAV-based GWAS encompassed the flowering time and photoperiod sensitivity data previously analysed by Bellucci et al. [13]. GWA analysis was run by using both the Mixed Linear Model (MLM) [67] and the Fixed and random model Circulating Probability Unification (FarmCPU) [68] implemented in the R package GAPIT v3 [69]. The threshold for each Genome Wide Association (GWA) scan was determined by the Bonferroni corrected *p* value at α = 0.05. The kinship matrix (IBS method) was calculated with Tassel 5 [70] and the population structure (at K2 obtained from Bellucci et al. [13]) were included into the models as fixed factors. Quantile-quantile (Q-Q) plots were obtained by plotting the observed -log10(*p* values) against the expected -log10(*p* values) under the null hypothesis of no association.

## Data Availability

The 109 raw sequencing reads generated and analyzed in this study have been deposited in the Sequence Read Archive (SRA) of the National Center of Biotechnology Information (NCBI) under BioProject number PRJNA1042929. Additional data comprising 230 raw sequencing reads have been sourced from Frascarelli et al. [11] and Bellucci et al. [13]. The pan-genome assembly and its annotation are publicly accessible via this link: https://doi.org/10.6084/m9.figshare.24573874.

